# Application of Blood Glucose and Ketone Meters for Measuring Glucose and Ketone Levels in Urine Samples

**DOI:** 10.1101/2025.09.03.674019

**Authors:** Zahra Panahi, Elizabeth Denley, Edgar D. Goluch

## Abstract

**BACKGROUND:** Urine glucose dipstick tests have low sensitivity, particularly in detecting glucose in early or mild cases of diabetes in cats and dogs. Ketone dipstick tests only detect acetoacetate and do not measure β-hydroxybutyrate (BHB) which is the predominant ketone in diabetic patients and the most reliable indicator of diabetic ketoacidosis (DKA).

**OBJECTIVE:** To evaluate the accuracy, sensitivity, and specificity of urine dipstick tests for glucose and ketone measurements in cats and dogs. To assess whether blood-based Fora meters provide more reliable urine measurements for diabetes detection.

**METHODS:** Standard solutions of glucose, BHB, and acetoacetate in artificial urine were analyzed using Fora meters and dipstick tests. Urine samples from 37 random dogs and 27 random cats were tested and results were analyzed using regression analysis, confusion matrices, and Bland-Altman plots.

**RESULTS:** Fora meters accurately detected glucose and BHP ketone levels in artificial urine with strong linearity to standards. Urine dipstick tests showed moderate accuracy (74%) but low sensitivity (57%) for glucose; performance varied by species and specific gravity. Urine dipstick test underestimate specific gravity in samples with high pH or high glucose levels. Urine dipstick tests cannot be used for BHP ketone measurements (sensitivity < 1%).

**CLINICAL RELEVANCE:** Fora meters even though originally designed for blood, may offer a more precise, point-of-care alternative to dipsticks for diagnosing diabetes and DKA in cats and dogs. Improved urine glucose and ketone quantification can help reduce false diagnosis and support earlier and more accurate treatment decisions.

## 1. Introduction

Diabetes in dogs is usually an autoimmune disease resembling Type 1 diabetes in humans and the most common cause of diabetes in dogs is destruction of insulin producing beta cells in the pancreas by immune system attacks. However, similar to Type 2 diabetes in humans, obesity, pancreatitis (inflammation of the pancreas) and pancreatic amyloid deposition (buildup of protein in pancreas) are major risk factors of diabetes in cats (1,2).

Measurement of fasting hyperglycemia (fasting blood glucose) and glucosuria (glucose in urine) are standard tests for diagnosing diabetes in dogs and cats. However, diagnosing diabetes in cats is more challenging than dogs since cats, on one hand, can show temporary elevated blood glucose levels due to stress leading to a false-positive diagnosis, and on the other hand, have a higher renal threshold for glucose (289 mg/dL for cats compared to 180 mg/dL for dogs (2)), allowing their kidneys to tolerate elevated blood sugar levels longer before glucose appears in the urine, which may result in a false-negative diagnosis (3). Additionally, many laboratories use dipstick tests to measure urine glucose levels and screen for diabetes. However, while urine dipstick readings for dogs and cats are useful for confirming glucosuria when positive, their low sensitivity makes them unreliable for ruling it out when negative (4).

Diabetic ketoacidosis (DKA), a life-threatening complication of diabetes, occurs when the body cannot utilize glucose for energy due to insulin deficiency. As a result, lipolysis in adipose tissue releases free fatty acids, which undergo ketogenesis in the liver, producing three main ketone byproducts: beta-hydroxybutyrate (BHB), acetoacetate, and acetone (5–7). While there are no commonly available point-of-care (POC) methods for measuring acetoacetate and acetone in blood (6), blood BHB can be measured using both laboratory tests and POC devices (8,9). In one study, a blood BHP level >3 mmol/L (31.23 mg/dL) was used as the diagnostic cut off for DKA in dogs and cats (10). Another study reported blood BHP level >3.8 mmol/L (39.56 mg/dL) as the most reliable indicator of DKA in dogs (11). For cats, blood ketone levels above 2.4-2.5mmol/L (24.99-26.03 mg/dL) were used as a diagnostic indicator of DKA. Values below these thresholds especially those under 1– 2 mmol/L (10.41-20.82 mg/dL) suggest that while the patient may be ketotic, ketosis is unlikely to be the primary cause of systemic condition (12,13). However, in urine, healthy patients should not have any detectable ketones, as they are fully reabsorbed by the renal proximal tubules (7). The urine dipstick tests and nitroprusside-based tablets are the most commonly used methods for detecting ketones in cats and dogs; however, these tests primarily detect acetoacetate and to a lesser extent, acetone, while they do not measure BHB (7–9). Since BHB is the dominant ketone in diabetes, these tests often tend to underestimate ketone levels by not detecting BHB. Additionally, false negatives may occur due to delayed sampling as acetone evaporates easily and acetoacetate can degrade due to bacterial activity (7).

The goal of this study is to assess whether measurement of urine glucose and ketone concentrations using glucose and ketone meters, originally designed for blood measurements, offer a more precise alternative for diagnosing diabetes in cats and dogs. To achieve this, we first determine the meters’ sensitivity in detecting standard concentrations of glucose, BHB, acetoacetate, and acetone in artificial urine solutions. We then compare their performance to that of urine dipstick tests, providing a comprehensive assessment of their diagnostic reliability.

## 2. Materials and Methods

### 2.1 Materials

Dextrose (D-glucose) anhydrous (Fisher Chemical, Certified ACS grade), sodium 3-hydroxybutyrate (Thermo Scientific Chemicals, 98%), lithium acetoacetate (Sigma-Aldrich, ≥90%), and artificial urine medium (Pickering Laboratories) were purchased from different suppliers and used in this study.

### 2.2 Animals and Sampling

Urine samples used in this study included 37 randomly selected dogs and 28 randomly selected cats from QSM Diagnostics patients. Pet owners conveniently collected these urine samples from their cats and dogs using FetchDx mail-in diagnostic testing kits at home and submitted to QSM Diagnostics Inc. for complete urinalysis from December 2024 to February 2025.

### 2.3 Measurements Using Fora Glucose and Ketone Meters

Different glucose and BHP ketone levels were measured using Fora test strips together with Fora 6 Connect instrument. To evaluate whether Fora test strips, originally designed for blood-based measurements, can be used to determine analyte concentrations in urine, standard solutions were prepared and tested. Standard solutions of glucose, BHB, and acetoacetate were prepared in artificial urine. These solutions were then tested at room temperature using the Fora meter to assess its performance across different matrices. Additionally, glucose and ketone (BHP) levels in 37 dogs and 27 cats’ urine samples were measured using the Fora test strips and Fora 6 Connect meter.

### 2.4 Measurements Using Insight Urinalysis Test Strips

Standard solutions of glucose, BHB, and acetoacetate in artificial urine were tested using InSight urinalysis veterinary urine test strips in conjunction with the Insight urinalysis veterinary urine analyzer. Similarly, 37 dogs and 27 cats’ urine samples were analyzed to measure glucose, ketone (acetoacetate), and specific gravity.

### 2.5 Measurement of Specific Gravity Using Refractometer

Specific gravity of urine samples was measured using a veterinary handheld refractometer with separate scales for cats and dogs. A small volume of urine was placed on the refractometer’s prism and the specific gravity was determined by reading the appropriate scale for the respective species.

### 2.6 Statistical Analysis

All statistical analyses were performed using MATLAB R2023b. Method comparison studies in this work were performed using scatter plots with regression analysis, confusion metrics, and Bland-Altman plots.

To evaluate the accuracy of Fora test strips in urine, scatter plots were generated comparing known standard concentrations of glucose and BHB in artificial urine with the corresponding measurements obtained from the Fora meter. A linear regression analysis was performed for each analyte to assess how closely the test strip readings aligned with the true concentrations. Error bars represent standard deviations from three independent measurements (N=3). Bar graphs were used to plot the Insight glucose dipstick test results because the readings from the dipstick tests are not continuous values but rather fall into discrete levels.

Confusion metrics including accuracy, sensitivity, and specificity were used to evaluate the diagnostic performance of urine dipstick tests against reference glucose and ketone meter readings. Binary classification threshold was set at 100 mg/dL to differentiate between “High” and “Normal” glucose levels. Additionally, subgroup analyses were conducted by stratifying samples by species (cats vs. dogs) and specific gravity levels (<1.040 vs. ≥1.040) to explore potential confounders affecting glucose test accuracy. Confusion metrics were used to compare the performance of dipstick tests to predict glucose in samples with high and low specific gravity. Glucose binary classification thresholds were set to 100 and 22 mg/dL for high (≥1.040) and low (<1.040) specific gravity levels.

Bland-Altman analysis was used to assess agreement between refractometer vs. dipstick for measurement of specific gravity. For consistency, all refractometer values above 1.060 were rounded to 1.070. The analysis included calculations of mean bias and 95% limits of agreement. Separate Bland-Altman plots were generated for cats and dogs’ samples to assess method agreement across species. To explore potential effect of pH (low <6, medium 6–7, high >7) and glucose levels (low glucose<100mg/dL, high ≥100mg/dL) on specific gravity boxplots were generated to visualize distribution differences across groups and groups with p-value < 0.05 were considered statistically significant. and 5 mg/dL for BHP ketone.

Finally, a confusion matrix was plotted to evaluate the performance of the dipstick test (which measures acetoacetate) against reference ketone meter readings (which measures BHP). Binary classification threshold was set at 5 mg/dL to differentiate between “High” and “Normal” ketone levels.

## 3. Results and Discussion

The first phase of this study aims to validate whether Fora test strips, originally designed for blood-based measurements, can accurately determine target analyte concentrations in urine. Standard solutions of glucose, BHB, and acetoacetate were prepared in artificial urine and tested using Fora glucose test strips (for glucose) and Fora ketone test strips (for BHB and acetoacetate). **Figure 1A** shows the Fora glucose meter readings plotted against the actual glucose concentrations in artificial urine, ranging from 12.5 to 500 mg/dL. The strong linear relationship with a slight overestimation indicates excellent agreement between measured and actual values suggesting that the Fora glucose meter can reliably estimate glucose levels in urine. For comparison, the same glucose solutions were tested using the Insight urine dipstick test, and **Figure 1B** displays the corresponding glucose strip readings. Unlike the Fora glucose meter, the Insight dipstick test produced discrete range-based values. Due to their semi-quantitative nature, dipstick tests are not suitable for precise quantitative measurements.

**Figure 1.**
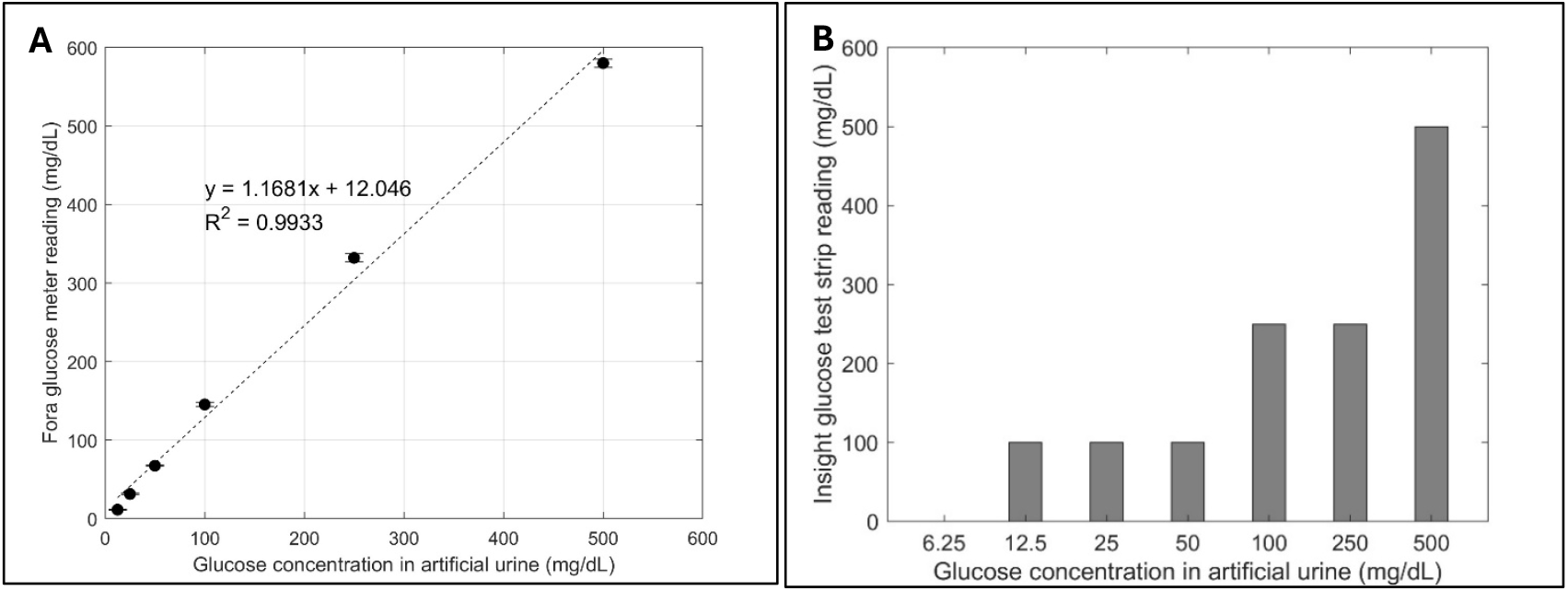
Glucose tests in artificial urine: **(A)** Fora glucose meter readings plotted against standard glucose concentrations in artificial urine. **(B)** Insight glucose dipstick test readings for the same glucose standards.

To evaluate the performance of the Fora ketone test for BHB detection in urine, standard solutions of BHB were prepared in artificial urine and tested using the Fora ketone meter. **Figure 2A** shows the Fora ketone meter readings plotted against BHB concentrations ranging from 6.25 to 80 mg/dL. The results demonstrate a strong linear correlation with a slight underestimation, suggesting that the Fora ketone meter can accurately quantify BHB in urine across a clinically relevant range. A 1000 mg/dL BHB solution tested with the Insight ketone dipstick test returned a negative result indicating its inability to detect BHB. In contrast, the Insight ketone dipstick test was used to detect standard acetoacetate solutions in artificial urine as shown in **Figure 2B**. This result suggests that ketone dipstick test can be used for semi-quantitative and broad screening of acetoacetate in urine. A 330 mg/dL acetoacetate solution tested with the Fora ketone meter produced no detectable signal confirming that this meter is not suitable for acetoacetate detection. These findings indicate that the Fora ketone meter is reliable for quantitative BHB measurement in urine, while the Insight dipstick is better suited for broad, semi-quantitative screening of acetoacetate.

**Figure 2.**
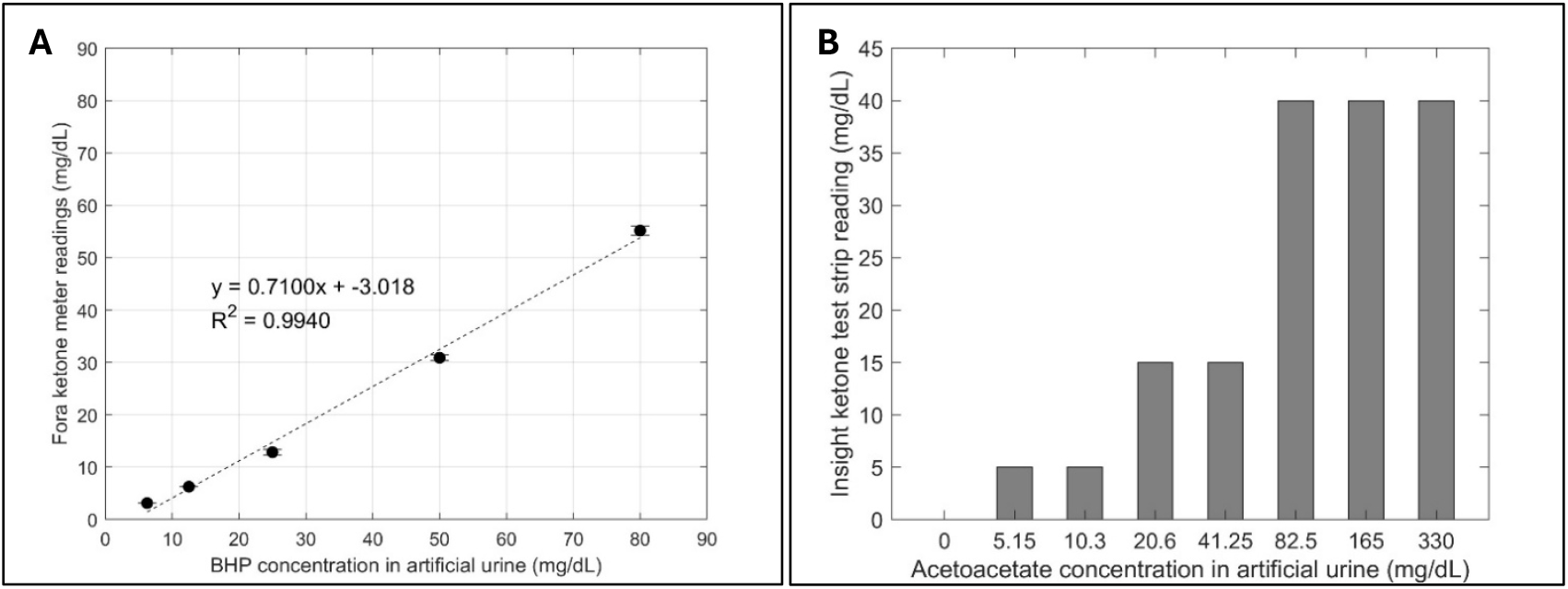
Ketone tests in artificial urine: **(A)** Fora ketone meter readings plotted against standard BHP concentrations in artificial urine. **(B)** Insight ketone dipstick test readings for standard acetoacetate concentrations in artificial urine.

The Insight glucose test strip is based on the specific glucose oxidase/peroxidase reaction, where glucose oxidase converts urine’s glucose into gluconic acid and hydrogen peroxide. Peroxidase on the test strip then reacts with hydrogen peroxide to produce a color change indicating glucose concentration. According to the Insight dipstick test instructions, high ketone concentrations may cause false negatives for samples containing small amounts of glucose (5.5 mmol/L or 99 mg/dl). Additionally, false positives may be caused by cleaning agents containing hypochlorite or peroxide. To assess the accuracy of the dipstick test in predicting glucose levels in urine, we compared its results to those from a glucose meter. **Figure 3A** shows the confusion matrix of Insight glucose dipstick test with reference to glucose meter readings for 65 urine samples from 28 cats and 37 dogs, using a cutoff of <100 mg/dL as “Normal” and ≥100 mg/dL as “High”. According to the confusion matrix, urine dip stick test can predict glucose levels with 73 % accuracy, 81% specificity and only 57% sensitivity. **Figure 3B** and **3C** show separate confusion matrices for cat and dog urine samples, respectively. The test predicted glucose levels in dog samples with slightly higher accuracy (76%) compared to cat samples (71%). Sensitivity was also higher in dogs (60%) than cats (56%), while specificity was greater in cat samples (92%) than in dog samples (78%). These findings suggest that while false negative readings are more common in cat samples, false positives are more frequent in dog samples.

**Figure 3.**
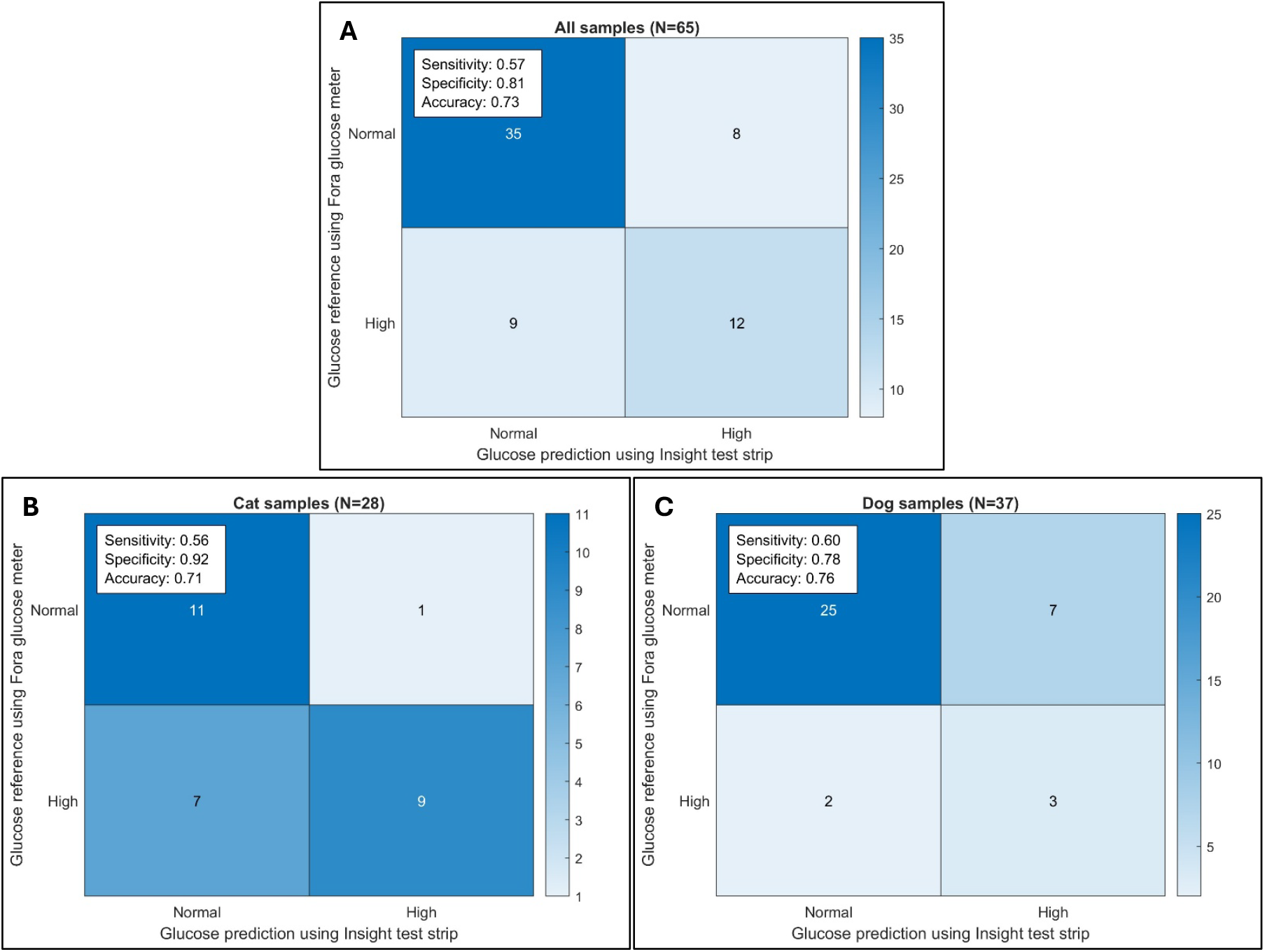
Confusion matrices evaluating the performance of the Insight glucose dipstick test against reference readings from the Fora glucose meter: **(A)** All urine samples (N=65). **(B)** Subset of cat urine samples (N=28). **(C)** Subset of dog urine samples (N=37).

To better understand the nature of false-negative and false-positive results, we examined the relationship between urine glucose levels and specific gravity to assess whether sample concentration influences test performance. **Figure 4** displays box plots comparing urine glucose levels measured by Fora glucose meter (**Figure 4A**) and Insight dipstick test (**Figure 4B**) across two specific gravity groups: below 1.040 (N=36) and 1.040 or higher than 1.040 (N=29). Specific gravity was determined using a refractometer. Both Fora glucose meter and Insight dipstick test demonstrated significantly elevated glucose levels in samples with higher specific gravity (P-value=9.8 × 10^−16^ for glucose meter, P-value=7.54× 10^−4^ for dipstick). This observation aligns with previous research indicating that increased urine glucose levels, contributes to higher specific gravity, especially in less concentrated urine (14).

**Figure 4.**
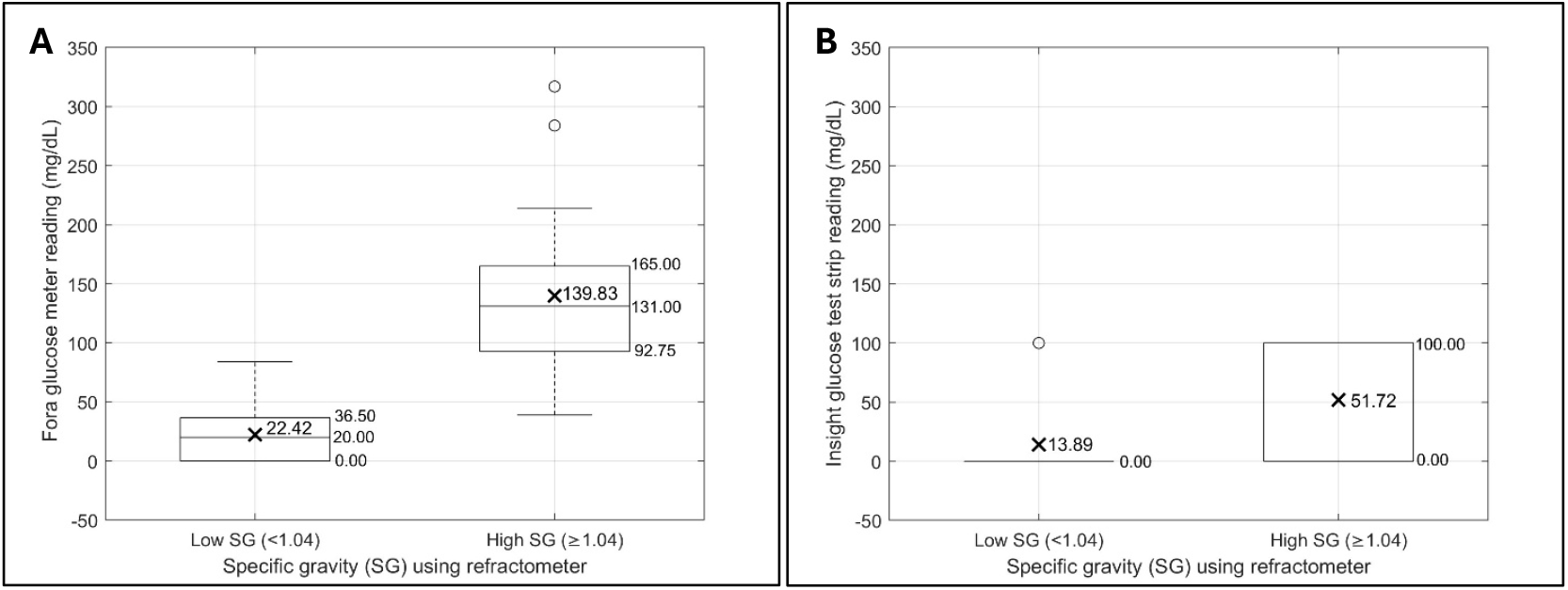
Comparison of urine glucose levels across low and high specific gravity groups measured using **(A)** Fora glucose meter and **(B)** Insight glucose dipstick test.

According to the Insight dipstick test instructions, reactivity of the glucose test decreases as the specific gravity of the urine increases. This means that the test is expected to be less accurate at higher specific gravities. To investigate the clinical implications of this, we evaluated the dipstick’s accuracy in predicting glucose levels in high (**Figure 5A**) and low (**Figure 5B**) specific gravity samples. A lower glucose cut-off of 22 mg/dL was applied to low specific gravity samples (below 22 mg/dL is “Normal” and above 22 mg/dL is “High”), reflecting their inherently lower glucose concentrations, while 100 mg/dL glucose cut-off was maintained for high specific gravity samples. Although the dipstick’s sensitivity was higher for high specific gravity samples (57% vs. 31%), its overall accuracy was lower (59% vs. 69%). The lower accuracy of high specific gravity samples is primarily due to a lower specificity (62%) compared to low specific gravity samples (100%).

**Figure 5.**
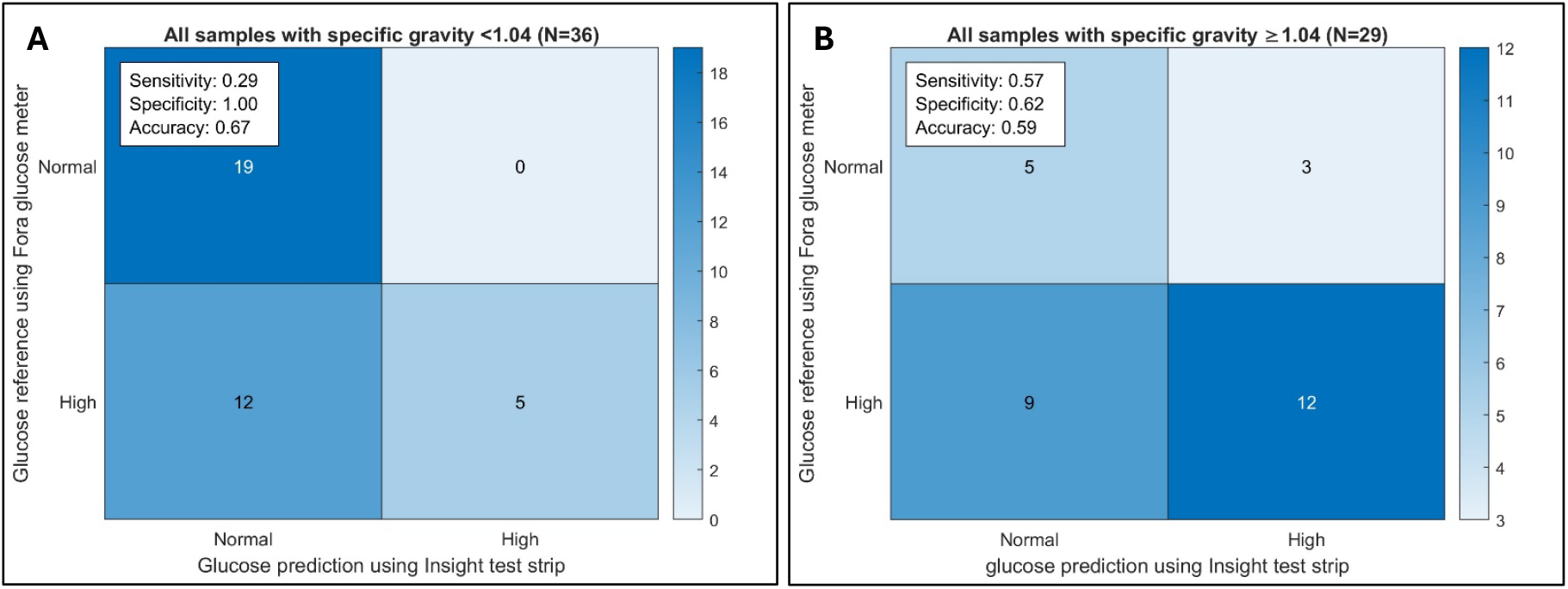
Confusion matrices evaluating the performance of the Insight glucose dipstick test against reference readings from the Fora glucose meter: **(A)** Low specific gravity samples (<1.040) (N=36) and **(B)** High specific gravity samples (≥1.040) (N=29).

We also evaluated the accuracy of the Insight dipstick test in measuring the specific gravity of urine samples. According to the test instructions, the strip contains a detergent and bromothymol blue which changes color from green to yellow in response to ionic constituents in urine. Specific gravity of all urine samples was also measured using a refractometer which permits more precise measurements across a wider range. The Bland-Altman analysis in **Figure 6A** shows the 95% limits of agreement between the dipstick and refractometer measurements ranging from 1.000 to 1.070. Overall, there was good agreement between the two methods with a small negative bias of 0.003, indicating that the dipstick tends to slightly underestimate specific gravity compared to the refractometer. However, the standard deviation increased at higher specific gravities suggesting greater variability and reduced reliability of Insight dipstick measurements in more concentrated samples. It should be mentioned that specific gravity dipstick tests are generally not recommended for cat urinalysis as urine specific gravity values are falsely decreased when urine pH exceeds 6.5 or when glucose is present in urine (7). Therefore, we also showed separate Bland-Altman plots for cats (**Figure 6A**) and dogs (**Figure 6B**). While dipstick readings for dog samples showed no significant bias, the dipstick slightly underestimated specific gravity in cat samples.

**Figure 6.**
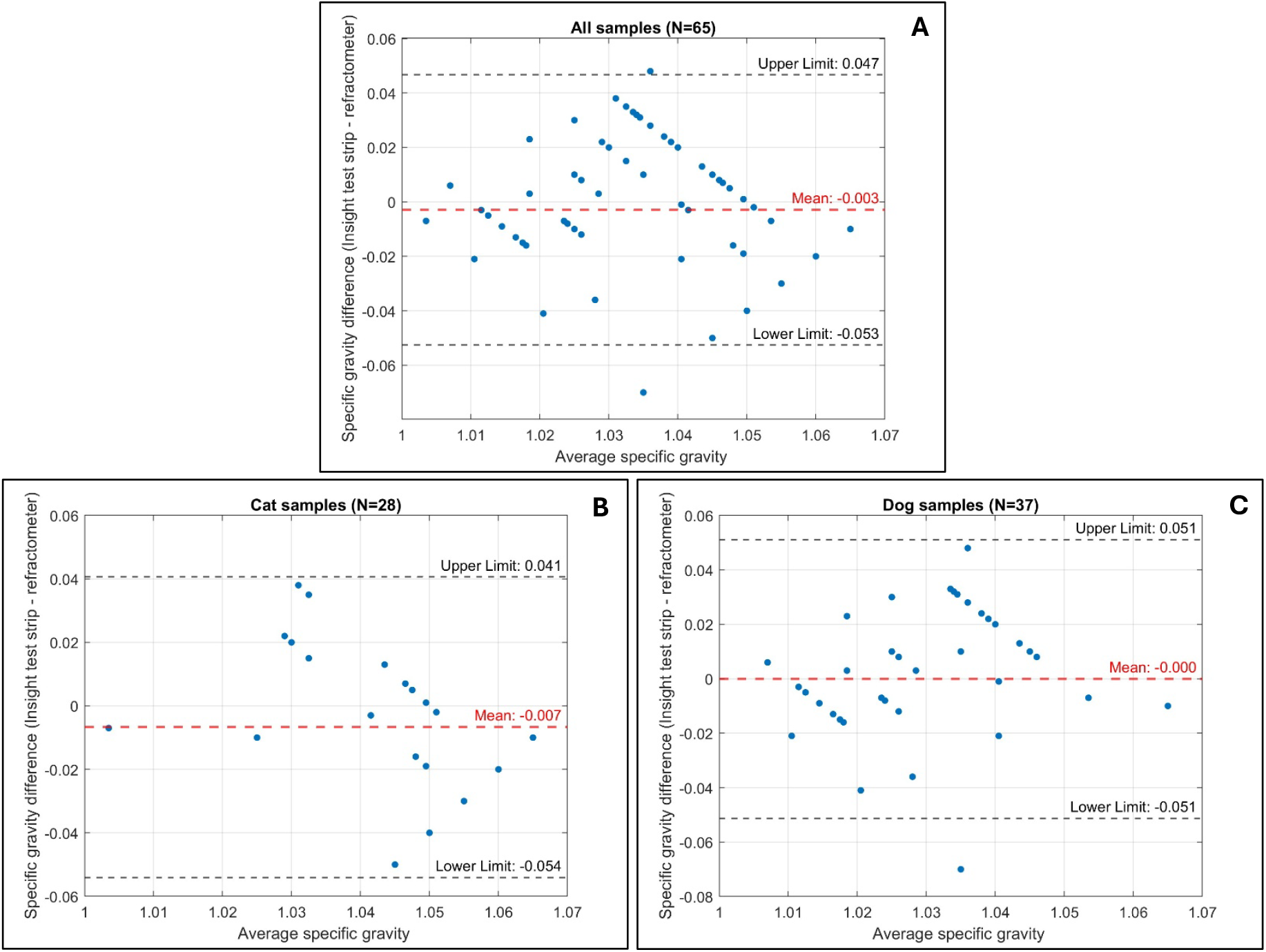
Bland-Altman plots evaluating the performance of the Insight specific gravity dipstick test against reference readings from refractometer: **(A)** All urine samples (N=65). **(B)** Subset of cat urine samples (N=28). **(C)** Subset of dog urine samples (N=37).

To assess whether urine pH affects the accuracy of specific gravity measurements, we analyzed urine samples across different pH ranges. We plotted urine specific gravity values measured using a refractometer (**Figure 7A**) and the Insight dipstick test (**Figure 7B**) for samples with low (pH <6), medium (pH 6–7), and high (pH >7) values. Highly buffered alkaline urine can lead to underestimation of specific gravity when using certain methods. However, according to the Insight dipstick test instructions, the analyzer automatically adjusts for pH effects when the urine pH is ≥7.0. However, our results show that while there is no statistically significant difference in refractometer-measured specific gravity among the pH groups (**Figure 7A**), the dipstick test reported significantly higher specific gravity values for high-pH samples compared to those with low or medium pH (**Figure 7B**).

**Figure 7.**
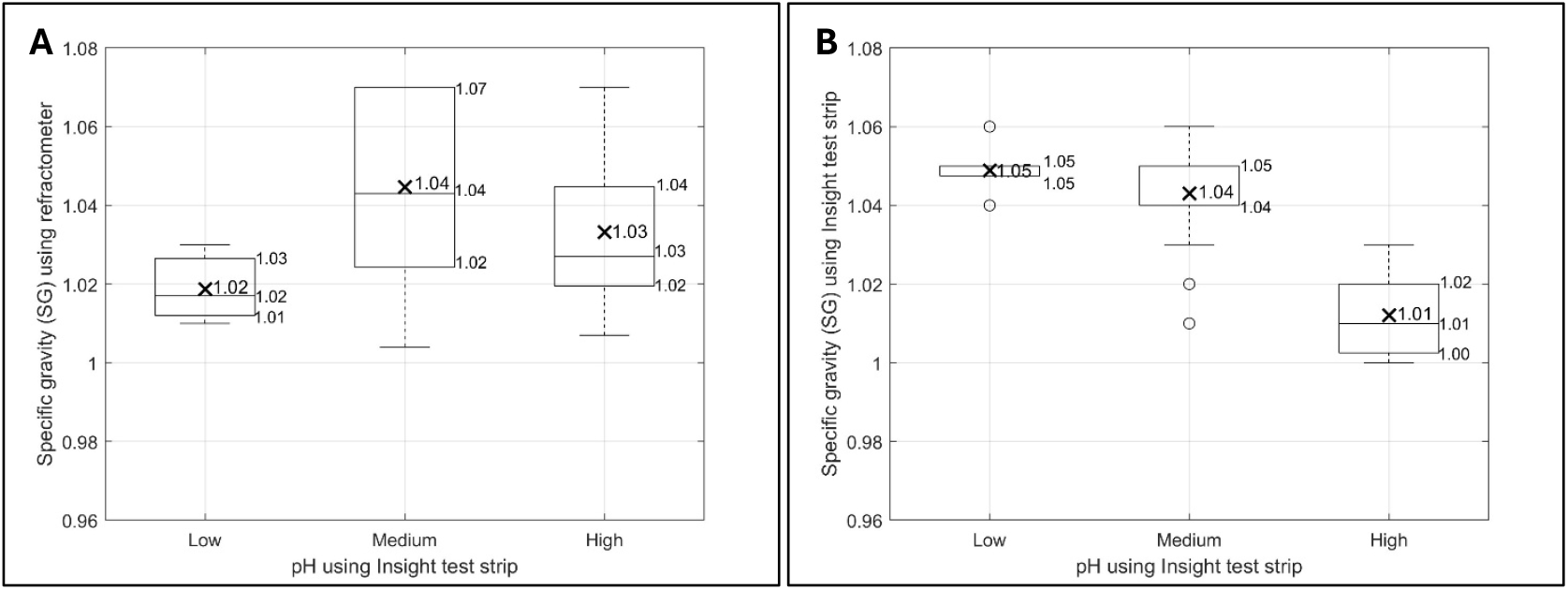
Comparison of urine specific gravities across low (pH <6), medium (pH 6–7), and high (pH >7) groups measured using **(A)** refractometer and **(B)** Insight specific gravity dipstick test.

To evaluate whether glucose levels in urine affect the accuracy of specific gravity measurements, we analyzed urine samples across different glucose concentrations. **Figure 8** shows specific gravity values measured using a refractometer (**Figure 8A**) and the Insight dipstick test (**Figure 8B**) for samples with low (<100 mg/dL) and high (≥100 mg/dL) glucose levels. While high-glucose samples had significantly higher specific gravity when measured by refractometer, there was no significant difference between low-glucose and high-glucose samples when measured by the dipstick test. These findings support the Insight test strip instructions which state that the dipstick may underestimate specific gravity in samples with high pH or high glucose levels.

**Figure 8.**
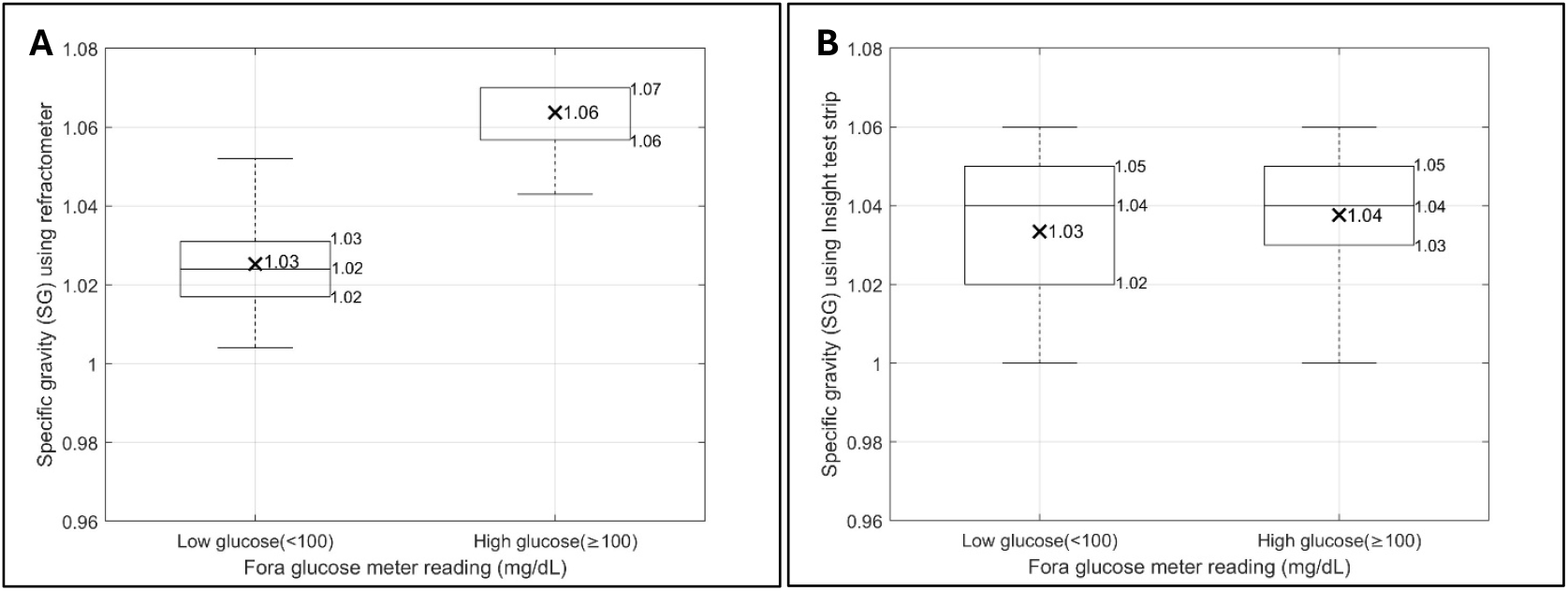
Comparison of urine specific gravities across low-glucose (<100 mg/dL) and high-glucose (≥100 mg/dL) groups measured using **(A)** refractometer and **(B)** Insight specific gravity dipstick test.

In the final part of this study, we evaluated the performance of the Insight dipstick test for detecting BHB in urine. **Figure 9** shows the confusion matrix comparing the Insight ketone dipstick test to the reference ketone meter readings for 65 urine samples. Since the dipstick test does not react with BHB and is only sensitive to acetoacetate, we expected no detection of BHB with this method. Consistent with this expectation, the dipstick identified only 2 out of 30 samples with high BHB levels (≥5 mg/dL). These two positive results may be explained by the presence of both BHB and acetoacetate in the samples since the test would detect acetoacetate. Alternatively, false positives may have occurred due to urine discoloration as the test instructions note that trace positive results can be caused by highly pigmented urine.

**Figure 9.**
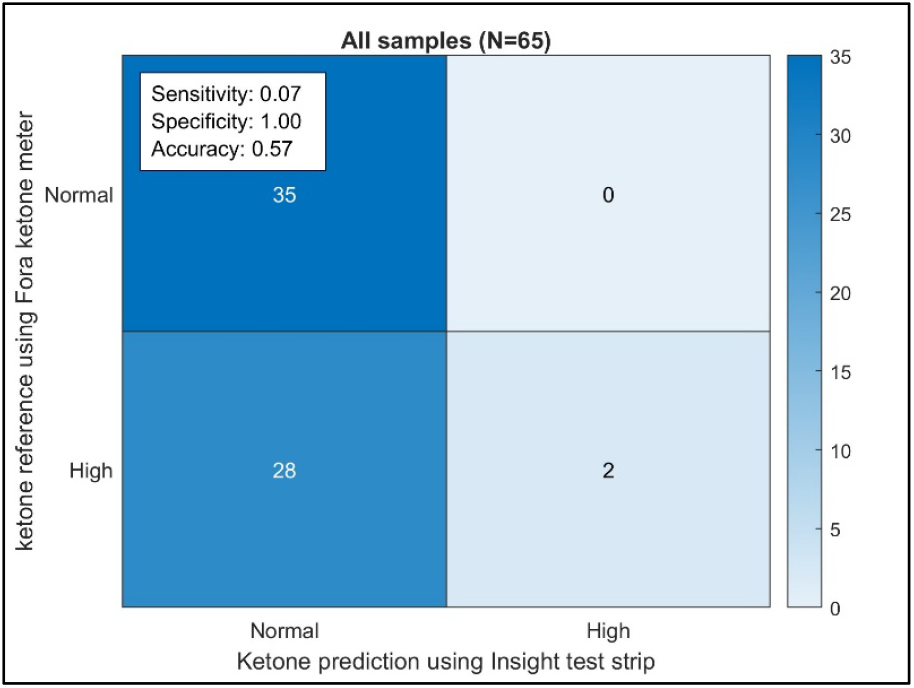
Confusion matrix evaluating the performance of the Insight ketone dipstick test against reference readings from the Fora ketone meter for all urine samples (N=65).

In summary, our findings demonstrate that Fora glucose and ketone meters offer a reliable and quantitative method for measuring glucose and BHB levels in urine. These meters showed excellent linear correlations with known standards. In contrast, glucose dipstick tests showed limited sensitivity, particularly in low-glucose and high-specific gravity samples, and were unable to detect BHB, the primary ketone associated with diabetic ketoacidosis. Given the semi-quantitative nature and susceptibility of dipsticks to different confounding factors, Fora meters provide a more robust alternative for early detection of diabetes in veterinary settings.

